# Unsupervised alignment reveals shared but locally refined colour qualia structure from toddlerhood to adulthood

**DOI:** 10.64898/2026.09.14.751604

**Authors:** Ryoichi Watanabe, Jue Wang, Masafumi Oizumi, Noburo Saji, Yusuke Moriguchi

## Abstract

Whether subjective colour experience (e.g., experience of ‘red’) is shared across children and adults remains a foundational question in developmental psychology and consciousness research. To test this, we compared the relational structures of their colour experiences and identified the mapping that best preserved these structures. Using pairwise similarity judgements for 13 basic colours, we applied Gromov–Wasserstein optimal transport (GWOT)—an unsupervised method requiring no prespecified stimulus–experience correspondence—to compare toddlers, preschool children, and adults. Preschoolers’ and adults’ group-averaged structures perfectly aligned, whereas toddlers’ structure did not fully align with either older group. However, removing specific colours—particularly those with relatively late-acquired linguistic boundaries, such as light blue—restored near-perfect alignment. Moreover, toddlers with greater colour-word knowledge showed closer correspondence with the adult similarity structure. These findings suggest that the global relational geometry of colour experience is largely established by toddlerhood, with limited local differences linked to colour-language acquisition.

## Introduction

Children and adults inhabit the same world, encounter the same objects, and often use the same colour words. Yet even when both call a strawberry red, it does not follow that they experience that redness (or ‘red’) in the same manner. This developmental issue relates to the classic inverted-qualia problem: *is my ‘red’ the same as your ‘red’?*^1–3^, but here the question is whether and how subjective colour experience differs across development. Traditionally, such questions have been treated as scientifically inaccessible because qualia—the qualitative aspects of conscious experience—are private and difficult to articulate ^4–6^.

Recent methodological advances, however, offer a way to make empirical progress on this question by reframing it as a structure-matching problem ^2,7,8^. Rather than asking what redness ‘feels like’ in absolute terms, this approach focusses on the relational organisation of colour experience: how each colour is positioned within a structure of similarities and differences relative to other colours ^9–11^. In this framework, pairwise similarity judgements between colours are compiled into a dissimilarity matrix, from which the putative similarity structure is inferred and visualised, for example, via multidimensional scaling (MDS). The degree of correspondence between the similarity structures of different individuals or groups can then be assessed quantitatively ^10,12,13^. Using this approach, our recent work introduced a child-friendly similarity-judgement task and demonstrated that colour similarity structures in Japanese and Chinese children aged 3–12 years were broadly similar to those of Japanese adults when evaluated by the correlation between the dissimilarity matrices as in conventional representational similarity analysis (RSA)—a supervised comparison in the sense that it assumes a predefined one-to-one correspondence between the colours presented to children and adults (e.g., children’s ‘red’ corresponds to adults’ ‘red’) (Figure 1A ‘supervised comparison’)^14^. The finding suggested that the global colour similarity structure emerges surprisingly early.

**Figure 1.**
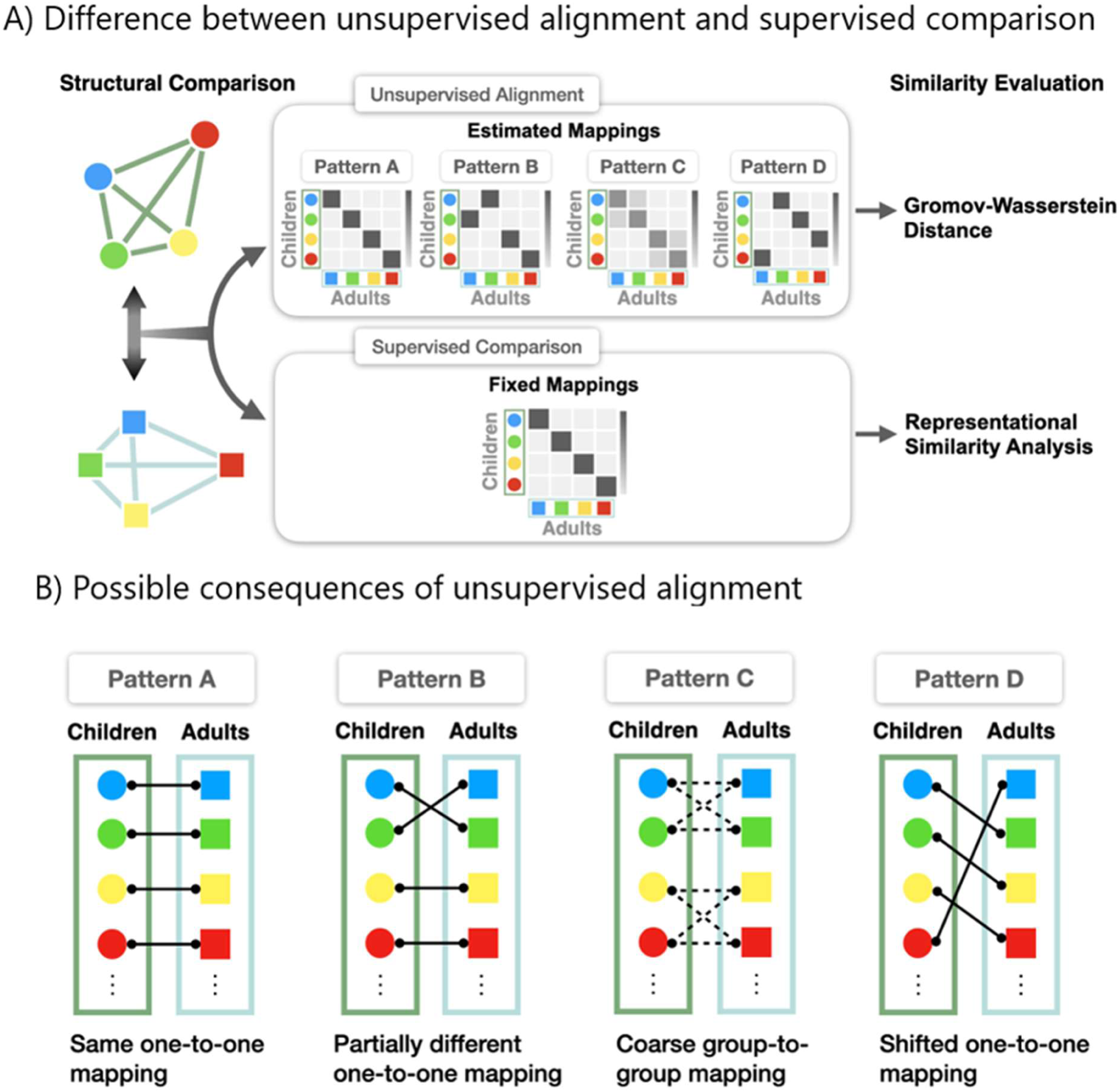
A) Differences between unsupervised alignment and supervised comparison. The former infers the optimal mapping from the relational structure itself, rather than assuming it from the outset and the latter evaluates the similarity between two representational structures under a single, prespecified correspondence between stimuli. B) Possible consequences of unsupervised alignment. Each pattern corresponds to each pattern in the section A. Pattern A indicates same one-to-one mapping, Pattern B indicates partially different one-to-one mapping, Pattern C indicates coarse group-to-group mapping, and Pattern D indicates shifted one-to-one mapping between children and adults.

This identity-based mapping in conventional RSA is a principled and often well-motivated choice ^12,15^. However, this assumption raises a question that RSA is not designed to answer: whether structurally equivalent spaces might also exist under alternative correspondences? This possibility is conceptually related to the inverted-qualia problem, but can be framed here as an empirical question about structural correspondence rather than as a direct test of phenomenological identity ^2^. For example, consider a simplified circular colour space consisting of four colours arranged in a circle: red, yellow, green, and blue. In the adult space, red is adjacent to yellow and blue, yellow is adjacent to red and green, and so on. Now suppose that children’s colour space has the same circular structure, but that the whole structure is shifted by one step relative to the adult structure. In this hypothetical case, the experience that children call ‘red’ would occupy the position that ‘yellow’ occupies in the adult space, children’s ‘yellow’ would occupy the position of adult ‘green’. Importantly, this kind of shift would preserve the relational pattern among colours. Adjacent colours would remain adjacent, and distant colours would remain distant, even if children’s individual colour experiences were not aligned with the same adult labels. Thus, children’s ‘red’ and ‘yellow’ could still be close to each other, just as adults’ ‘red’ and ‘yellow’ are close to each other, not because children’s ‘red’ necessarily corresponds to adults’ ‘red’, but because the overall circular structure has been preserved. Consequently, the overall pattern of pairwise dissimilarities could remain highly similar across children and adults, even though the best one-to-one correspondence between individual colour experiences would not be ‘red-to-red’, ‘yellow-to-yellow’, and so forth. A high RSA correlation would, therefore, indicate that the two spaces have similar relational organisation under the imposed label-based ordering, but it would leave open whether another mapping between the two spaces would provide an equally good, or even better, correspondence.

To address this possibility, an alignment method is required that infers the optimal mapping from the relational structure itself, rather than assuming it from the outset ^7,16^. Gromov–Wasserstein optimal transport (GWOT) provides exactly this capability ^7,17^. GWOT searches for a mapping between two spaces that preserves pairwise distance relation as much as possible. In this sense, it provides a data-driven correspondence between positions in two psychological colour spaces, rather than requiring the researcher to impose a ‘red-to-red’, ‘blue-to-blue’, and so forth correspondence in advance. The resulting mapping can then be evaluated by asking whether each colour is aligned with its adult counterpart—for example, whether children’s red is mapped to adults’ red—without this correspondence having been assumed when estimating the mapping (Figure 1A, B). Recent applications have shown that GWOT can robustly align colour similarity structures within colour-neurotypical adults, while revealing systematic misalignment between neurotypical and colour-atypical individuals ^7,16^. Applying this framework to developmental data, therefore, offers a more stringent test of structural equivalence than correlation-based analyses alone, because it asks not only whether children’s and adults’ colour spaces have similar relational organisation, but also whether the label-based correspondence itself is supported by the geometry of the data.

In this study, we apply this framework to compare younger children, older children, and adults. This developmental comparison is motivated by a long-standing debate over the role of language in shaping colour experience. Substantial evidence suggests that key aspects of colour organisation are biologically grounded and emerge before colour-word acquisition. Infants as young as four months show categorical perception of hue ^18–21^, and a large-scale mapping study identified five broad infant colour categories whose boundaries appeared to be partly constrained by early cone-opponent colour-vision mechanisms ^22^. These findings suggest that the basic structure of colour experience can emerge during early development. At the same time, language acquisition may affect the structure of colour experience. Indeed, several studies have shown that language can modulate aspects of colour processing, particularly around category boundaries ^23–29^. However, most previous studies have examined relatively local distinctions, such as blue–green or red–pink boundaries. Therefore, it remains vague whether language reshapes the global structure of colour experience, or whether its influence is largely confined to local discriminations and lexical category boundaries.

This question is especially important developmentally. Previous studies of children’s colour similarity structures have focussed on ages three years and above ^14^ by which age many children have already acquired much of the basic colour lexicon ^30–32^. Toddlers aged 2–3 years, by contrast, are still in the midst of active colour-word acquisition ^32–34^. They, therefore, provide a critical test of whether broad structural alignment with adults is already present before colour-term knowledge has fully stabilised. Preschool children provide a complementary comparison group, because their colour vocabulary is expected to be substantially more consolidated. Comparing these two child groups with adults enables testing whether developmental changes in colour-term knowledge are accompanied by changes in similarity structure.

Here, we investigate the colour similarity structures of toddlers, preschool children, and adults using pairwise similarity judgements for 13 colours corresponding to the basic Japanese colour terms (including light blue and light green). We apply GWOT to test whether the colour similarity structures of these groups can be aligned in a fully unsupervised manner. Additionally, we assess colour-word comprehension and production in the toddler group and examine whether individual differences in colour-term knowledge modulate the correspondence between toddlers’ and adults’ similarity structures at the trial level. If the global colour similarity structure is already in place by toddlerhood, we would expect robust alignment for the majority of colours, with misalignment confined to specific regions where linguistic boundaries are still being acquired. Conversely, if language fundamentally reorganises colour experience, we would expect pervasive structural differences between toddlers and older groups, and performances of colour-word tasks would modulate toddlers’ colour similarity structure.

## Results

### Colour similarity structures across age groups

We recruited 40 toddlers, 32 preschool children, and 32 adults, and administered a smartphone-based colour similarity task using touch responses (see Figure 2 and Methods). After applying our exclusion criteria based on task completion and double-pass reliability (*r* > 0.30) ^35^, 31 toddlers, 28 children, and 32 adults were retained for analysis. For preschool children and adults, each participant rated all 169 ordered colour-pair trials among 13 colours, including same-colour pairs on a 4-point scale (very similar, slightly similar, slightly dissimilar, very dissimilar). For toddlers, each participant rated a subset of 30 pairs on a 2-point scale (similar, dissimilar), and individual partial matrices were integrated at the group level to alleviate extraneous task demands. Previous work on children’s self-report and rating-scale use suggests that young children often experience difficulty using graded response categories reliably, tend to rely on extreme or dichotomous responses, and may struggle especially with intermediate scale points ^36,37^. Thus, the binary format was intended to maximise task comprehension and response reliability in toddlers, while preserving the core requirement of making pairwise similarity judgements. Given the difference in response scales between toddlers and preschoolers/adults, we conducted supplemental analyses to examine whether this scale difference affected the results. Specifically, the 4-point similarity ratings from preschoolers and adults were recoded into a binary scale corresponding to the toddler response format, and the main analyses were repeated using these recoded data. The results were consistent with those reported in the main text (Figure S1-2).

**Figure 2.**
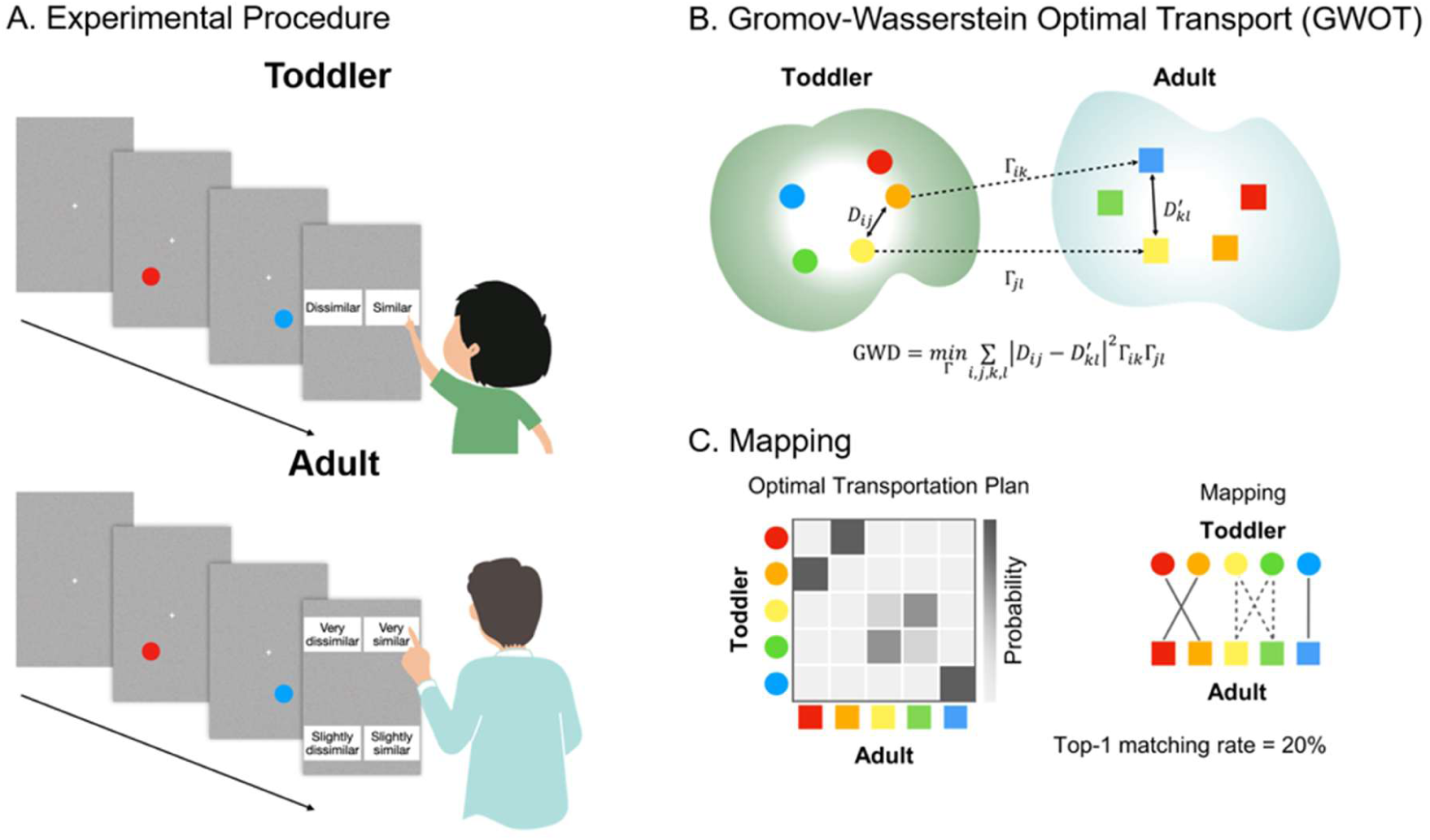
Overview of the study. A) Experimental procedure. To estimate the colour similarity structure for each group, toddlers, preschool children, and adults completed pairwise similarity judgements for 13 colours. Colour similarity structures are estimated for each group by applying multidimensional scaling (MDS). B) Gromov–Wasserstein optimal transport (GWOT). The elements of D and D′ are the distances (dissimilarities) between the colour experiences for each group. Γ is the transportation plan, which designates the correspondence between colour similarity structures. GWOT finds the optimal correspondence between two structures that preserves the distance relationship between structures as much as possible. C) Example of the optimal transportation plan Γ (left panel). The optimal transportation plan can be seen as a mapping between two structures (right panel).

Figure 3A illustrates the group-averaged dissimilarity matrices. Visual inspection reveals broadly similar patterns across all three groups. Two-dimensional MDS solutions confirm this overall organisation (Figure 3B): all three groups produce an approximate colour ring in which perceptually similar hues are proximal. However, the MDS solution for toddlers shows notable deviations, particularly in the arrangement of blue, light blue, and purple, which appear displaced relative to the corresponding positions in the older groups.

**Figure 3.**
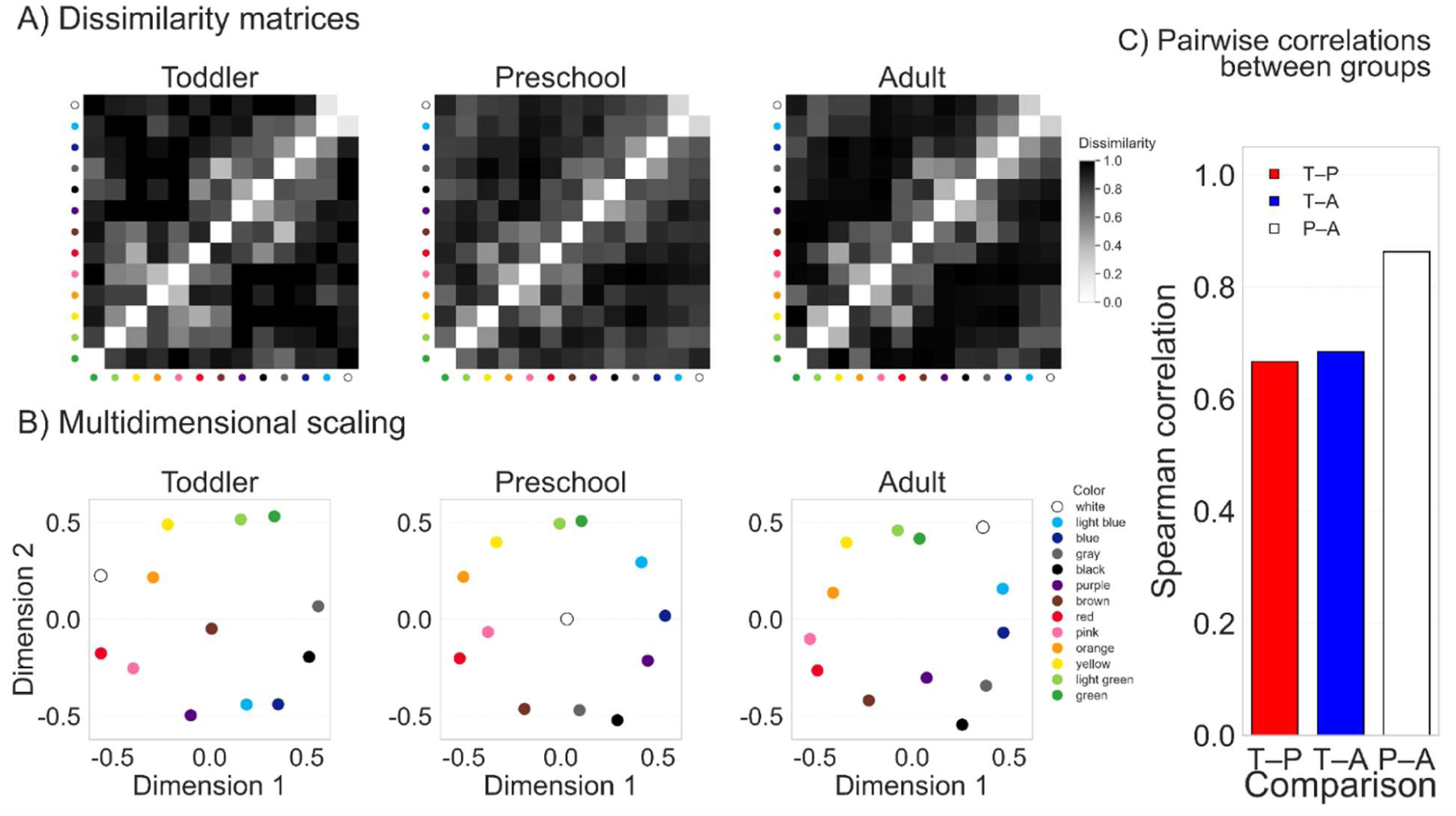
Analyses with the conventional RSA. A) Group mean dissimilarity matrices for each age group. White and dark cells represent high similarity and dissimilarity, respectively. B) 2-dimensional MDS representation in each age group. Symbol colours correspond to the colours used in the experiment. C) Pairwise correlations of dissimilarity matrices between groups. T-P indicates a comparison between toddlers and preschoolers, T-A indicates a comparison between toddlers and adults, and P-A indicates a comparison between preschoolers and adults.

To quantify these observations, we first conducted the conventional RSA. Specifically, we evaluated the correlation between the upper-triangular values of the group-level dissimilarity matrices using Spearman’s rank correlations, with significance assessed via permutation-based Mantel tests (5,000 permutations; Figure 3C). The correlation between preschool children and adults was high (ρ = 0.86, *p* < 0.001). In contrast, correlations involving toddlers were lower: ρ = 0.67 with preschool children and ρ = 0.69 with adults (*ps* < 0.001). These results confirm that the global pattern of colour similarity judgements is broadly preserved across all three groups, while also indicating divergence for the toddlers.

### Unsupervised alignment via GWOT

To identify the optimal mapping that best preserves structures between participant groups, we performed GWOT for each pair of age groups and evaluated alignment quality using the top-1 matching rate. It measures the probability that the experience of each colour in one participant group is mapped to that of the same colour in the other group under the optimal transportation plan (Figure 4A, B). Note that GWOT was performed without colour labels, but after the optimal mapping between colour experiences across different groups was identified by GWOT, colour labels were applied solely to evaluate the top-1 matching rate.

**Figure 4.**
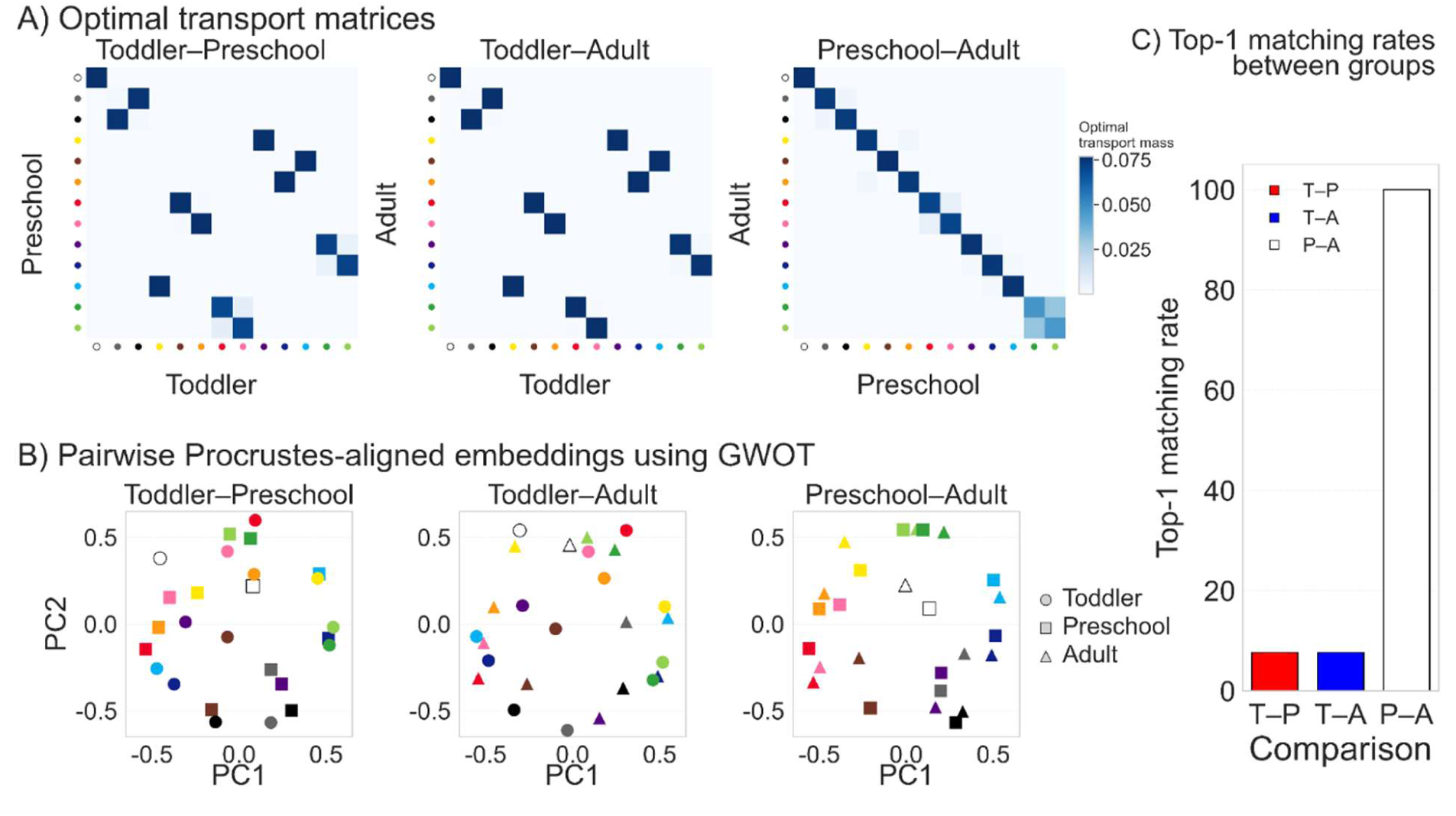
Analyses with GWOT. A) Optimal transportation matrices between groups. B) Pairwise Procrustes-aligned embeddings using GWOT. C) Top-1 matching rates between groups. T-P indicates a comparison between toddlers and preschoolers, T-A indicates a comparison between toddlers and adults, and P-A indicates a comparison between preschoolers and adults.

The alignment between preschool children and adults yielded a top-1 matching rate of 100% (Figure 4C), indicating that every colour of one group was mapped to its correct counterpart of the other group under purely unsupervised matching. This case corresponds to the same one-to-one mapping illustrated in Pattern A of Figure 1B. This perfect alignment demonstrates that the same-colour correspondence can be recovered from the relational structure itself, rather than specified in advance. The result extends our previous finding of high correlation to the stronger criterion of unsupervised recoverability ^14^. In contrast, the alignment between toddlers and preschool children yielded a top-1 matching rate of 7.7%, and that between toddlers and adults also yielded 7.7% (Figure 4C). These rates, at the chance level of 7.7% (1/13), indicate that the toddlers’ similarity structure deviates from those of older groups in such a manner that an exact one-to-one mapping does not preserve the structure optimally, resulting in a shifted one-to-one mapping.

### Leave-one-colour-out analysis reveals localised sources of misalignment

We reasoned that the failure of alignment between toddlers and the older groups could arise from either pervasive structural differences or localised deviations in specific colour regions. GWOT is sensitive to individual items that are anomalously positioned in one structure but not in the other: a single displaced colour (an outlier) can propagate misalignment across the entire transportation plan ^38^. Thus, if pervasive structural differences exist between toddlers and the older groups, removing any single colour should not substantially improve the matching rate. By contrast, if a localised deviation is responsible for the imperfect alignment, the matching rate should improve when the relevant colour is removed. To distinguish between these possibilities, we performed an exploratory partial structure matching analysis, called leave-one-colour-out analysis, in which each of the 13 colours was removed in turn and the GWOT alignment was recomputed on the remaining 12 colours.

The results revealed that the removal of specific colours markedly improved alignment scores (Figure 5A). In the comparison between toddlers and preschool children, the removal of light blue, white, black, purple, blue, and light green each yielded substantial improvements in the top-1 matching rate. In the comparison between toddlers and adults, the removal of light blue, white, black, yellow, and gray produced analogous improvements. Notably, light blue emerged as a consistent source of misalignment in both comparisons: removing light blue produced the largest single improvement, bringing the alignment score close to the preschool–adult level (Figure 5A), while also yielding the lower Gromov–Wasserstein distance (GWD; Figure S3A). Examination of the MDS solutions corroborates this finding: in the toddlers’ structure, light blue is positioned closer to purple, whereas in the preschool children and adult structures, it is positioned closer to green (Figure 3B). After light blue was removed, the MDS solutions showed near-perfect alignment (Figure S3B).

**Figure 5.**
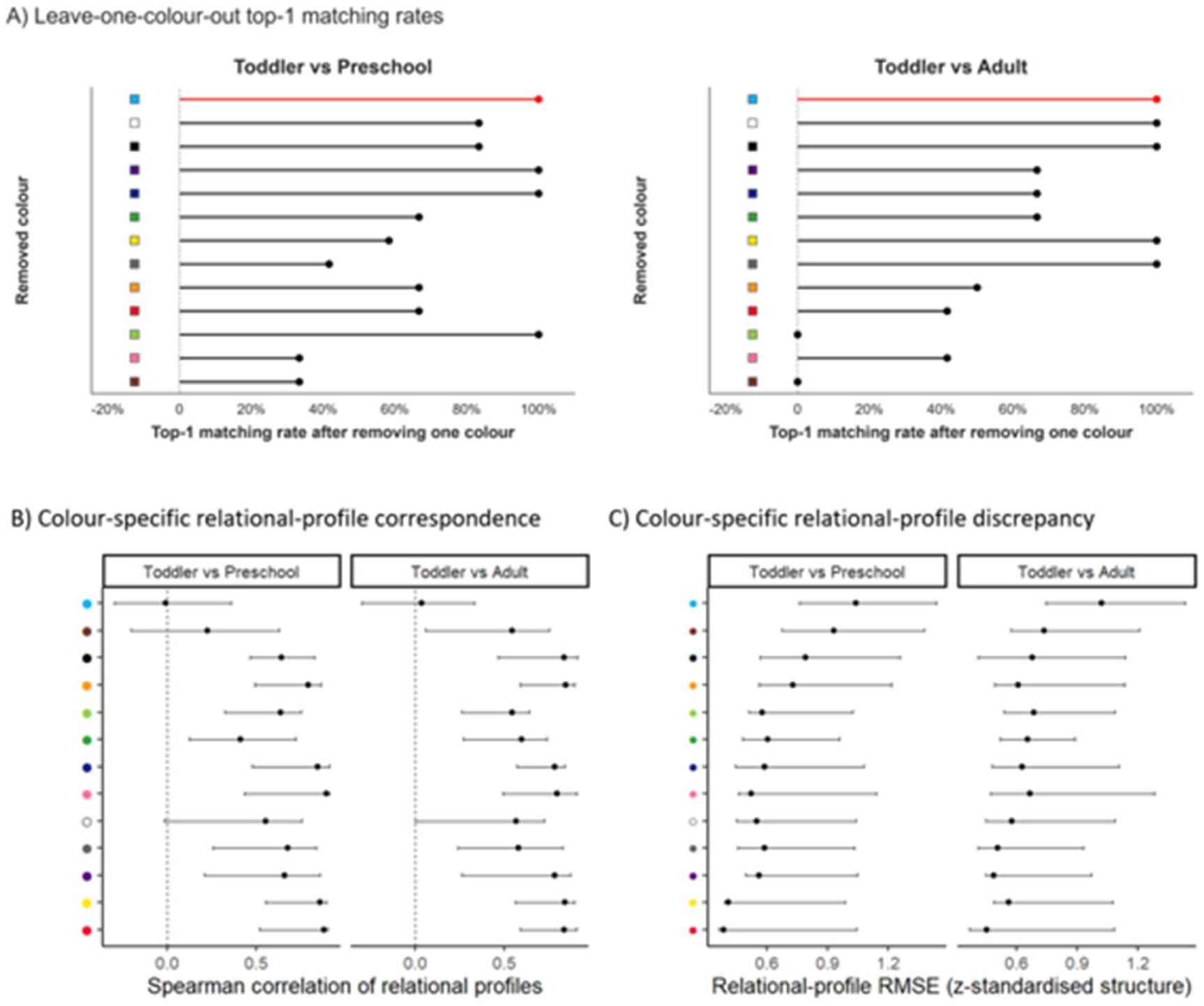
Colour-specific contributions to developmental differences in relational structure. A) Top-1 matching rates after removing each colour. B) Spearman correlations between toddler and preschool or adult relational profiles, each comprising a colour’s dissimilarities to the other 12 colours. Higher correlations indicate greater preservation of relational rank order. C) Standardised RMSEs between the corresponding profiles. Higher RMSEs indicate greater discrepancy. Error bars in B and C represent 95% Bayesian-bootstrap credible intervals.

To further assess whether the developmental differences were observed in the relational profile of a particular colour, especially light blue, or reflected a broader local structure centred on light blue, we conducted two complementary analyses. First, we estimated group-level dissimilarities for all 78 colour pairs and constructed, for each colour, a 12-element relational profile representing its dissimilarity to every other colour. Toddler profiles were compared with adult and preschool profiles using Spearman correlations and standardised RMSEs, with uncertainty estimated through participant-level Bayesian bootstrap resampling. Results revealed that light blue showed the clearest and most consistent developmental divergence (Figure 5B, C). Its relational profile was essentially uncorrelated with both the adult profile (ρ = .04, 95% Bayesian-bootstrap credible interval [−.305, .329]) and the preschool profile (ρ = −.01 [−.298, .361]), and it had the largest standardised RMSE in both comparisons (1.02 [.744, 1.435] and 1.04 [.761, 1.436], respectively). Other colours did not show comparable evidence of developmental divergence. For example, white, which also emerged as a potential contributor in the leave-one-colour-out analysis, showed moderate positive correspondence with both the adult profile (ρ = .57, 95% Bayesian-bootstrap interval [−.004, .725]) and the preschool profile (ρ = .56 [−.017, .760]), although the intervals narrowly included zero. Its standardised RMSEs were also substantially smaller than those for light blue in both comparisons (0.58 [.448, 1.086] and 0.55 [.447, 1.040], respectively). Thus, although some colours showed isolated signs of discrepancy in specific analyses, light blue was the only colour showing consistently weak profile correspondence and comparatively large profile-based discrepancies across both developmental comparisons.

Next, we examined whether the distinctive relational profile of light blue extended to a broader local substructure centred on it. We prespecified a four-colour subset comprising light blue, blue, white, and green based on adults’ MDS solutions, and compared the six pairwise dissimilarities within this subset across groups. The standardised local-structure RMSE was 0.687 [0.465, 1.210] for toddlers versus adults and 0.581 [0.394, 1.079] for toddlers versus preschool children. At the same time, the rank ordering of the six relations remained moderately to strongly preserved across groups (ρ = .714 [0.200, 0.886] and ρ = .943 [0.371, 1.000], respectively). Crucially, however, the light-blue-centred four-colour structure was not reliably more discrepant than equivalently sized four-colour subsets that comprised all the colours except light blue. To quantify this, we computed a matched-subset contrast by subtracting the mean RMSE of all four-colour subsets without light blue from the RMSE of the light-blue-centred subset; positive values therefore indicate greater discrepancy for the light-blue-centred subset than for comparable subsets elsewhere in the colour space. The standardised RMSE contrast was 0.147 [−0.262, 0.523] for toddlers versus adults and 0.043 [−0.348, 0.387] for toddlers versus preschool children. Sensitivity analyses using light-blue-centred subsets containing two to five colours yielded the same general pattern, with none of the corresponding contrast intervals excluding zero (Figure S4). Together, these findings indicate that developmental differences were most consistently evident in the relational profile of light blue and, particularly, relations involving it, but did not provide robust evidence that a broader light-blue-centred local structure was uniquely more discrepant than comparable substructures elsewhere in the colour space. These findings suggest that the global relational geometry of colour experience is broadly shared between toddlers and adults, with misalignment concentrated in a specific colour, light blue, whose Japanese lexical boundaries—notably the basic-level distinction between ao (blue) and mizuiro (light blue)—are acquired relatively late ^32^.

### Colour-word knowledge modulates alignment with the adult structure

Based on the leave-one-colour-out analysis, we examined whether the misalignment in toddlers’ similarity structures can reflect incomplete colour-word acquisition. To directly test this possibility, we assessed colour-word comprehension and production in the toddler group and examined their relationship to similarity-structure correspondence. One child was excluded from the analyses because of tasks incompletion. Mean correct performance was 82.8% (SD = 20.4) in the comprehension task, and mean correct performance was 70.0% (SD = 22.7) in the colour labelling task.

We examined whether individual differences in children’s colour-word knowledge relate to the correspondence between children’s and adults’ colour similarity structures. Specifically, we asked whether toddlers’ ‘similar’ responses for a given colour pair (e.g., a red–orange colour pair) track the adults’ ‘similar’ responses of that pair more strongly when the child has acquired the corresponding colour words (e.g., children acquired the colour words for ‘red’ and ‘orange’). For each child and each colour, we computed a colour-word knowledge score based on performance in the comprehension and labelling tasks, ranging from 0 to 2, given that performance on the two tasks was highly correlated. For each similarity-judgement trial involving colours a and b, we then defined a trial-level knowledge score as KnowSum_i(a,b) = KnowColour_i(a) + KnowColour_i(b), which ranges from 0 to 4. Higher KnowSum, therefore, indicates that the child has better knowledge of the two specific colour words involved in that trial (Figure 6A). To quantify the adult reference structure for each colour pair, we used the adult group-average dissimilarity values. For interpretability, we converted adult dissimilarity into an adult similarity anchor by reversing the scale, such that larger values indicate that adults judged the pair as more similar.

**Figure 6.**
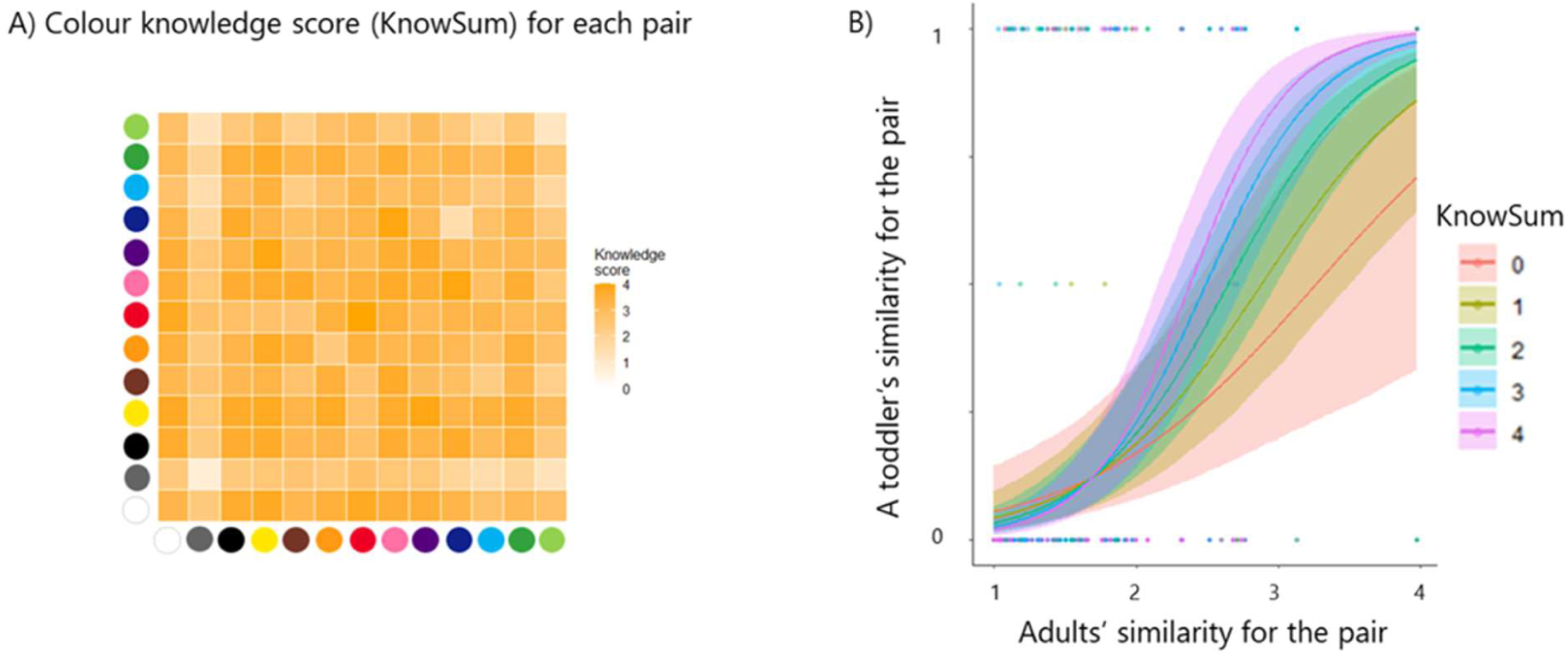
A) The correspondence between toddlers’ and adults’ ‘similar’ responses modulated by children’s colour word knowledge. B) Results. The y-axis represents predicted ‘similar’ responses for a given colour pair by toddlers, and the x-axis represents the adults’ ‘similar’ responses of that pair. KnowSum indicates that the child has better knowledge of the two specific colour words involved in that trial.

We fit a generalised linear mixed-effects model (GLMM) with a logit link to the toddlers’ binary similarity responses (similar/dissimilar). Predictors were mean-centred and included the adult similarity anchor (Adult_c; the reversed adult group-mean dissimilarity for each colour pair), a trial-level colour-word knowledge score (KnowSum_c; sum of the child’s comprehension and labelling scores for the two colours in the trial, ranging 0–4), and age (age_c). Random intercepts were included for participant and colour pair. The primary model was:

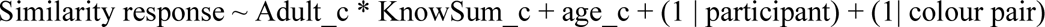

Results showed that the main effect of KnowSum_c was not significant (b = −0.015, SE = 0.113, z = -0.130, p = 0.896), but the Adult_c × KnowSum_c interaction was significant (b = 0.403, SE = 0.116, z = 3.464, p < 0.001; likelihood-ratio test: χ²(1) = 10.548, p = 0.001; Figure 6B). The positive interaction indicates that as toddlers’ knowledge of the two relevant colour words increased, their similarity judgements were more strongly correlated with the adults’ responses, suggesting that toddlers’ colour similarity structure can track the adult structure more closely with colour-term acquisition. Importantly, this effect was observed over and above the contribution of chronological age, suggesting that colour-word acquisition per se—rather than general maturation—is associated with the refinement of similarity-structure correspondence.

## Discussion

The present study applied an unsupervised alignment method, GWOT, to compare the colour similarity structures of toddlers, preschool children, and adults. Three main findings emerged. First, the colour similarity structures of preschool children and adults aligned perfectly under unsupervised matching (top-1 matching rate = 100%), demonstrating structural equivalence under a more stringent criterion than conventional correlation-based methods. Second, while toddlers failed to fully align with the older groups, a leave-one-colour-out analysis showed that near-perfect alignment was obtained when specific colours, such as light blue, was removed. Additional analyses identified light blue as the most consistently distinctive individual colour. This pattern suggests that the misalignment reflects differences in a limited configuration of relations involving light blue, rather than a pervasive difference across the entire colour space. Third, a trial-level analysis demonstrated that toddlers’ colour-word knowledge significantly modulated the degree to which their similarity judgements tracked the adult reference structure, independent of chronological age.

The perfect unsupervised alignment between preschool children and adults is, to our knowledge, the first demonstration that colour similarity structures can be aligned across developmental groups without relying on predefined colour labels or correspondences. This finding substantially extends the correlation-based evidence reported in our previous work ^14^ and is consistent with growing evidence from infant and toddler research that the core organisation of colour perception is biologically constrained and emerges early in development ^19–21^. Importantly, these perceptual foundations appear to be in place before colour-word acquisition ^22^. Our result implies that the relational geometry captured by these early categorical distinctions is preserved through childhood and into adulthood with remarkable fidelity, at least at the resolution of basic colour terms.

The finding that toddlers’ structures did not fully align with the older groups, yet showed near-perfect alignment upon removal of a specific colour, provides an important nuance. Rather than indicating a fundamentally different mode of colour experience in toddlerhood, the misalignment appears to arise from a restricted configuration of relations involving colours whose Japanese lexical boundaries are still being acquired. The most prominent source of misalignment was light blue (*mizuiro*)—a colour whose basic-colour-term status in Japanese ^39,40^ implies that children must learn to differentiate it from both blue (*ao*) and green (*midori*). Cross-linguistic studies have shown that the blue–light blue boundary is language-specific: Russian, which has obligatory basic terms for light blue (*goluboy*) and dark blue (*siniy*), produces enhanced discrimination at this boundary in adult speakers, whereas English speakers, who lack this distinction at the basic level, do not ^28^. The displacement of light blue in our toddler group—closer to purple and farther from green—is consistent with developmental evidence that younger children overextend ‘blue’ to stimuli adults would label ‘purple’ ^31,41^, and with the slow-mapping account of colour-word acquisition, which holds that children gradually refine lexical boundaries around preexisting perceptual categories ^31,32,42^.

The trial-level analysis provides complementary, individual-level evidence of the role of language. The significant Adult_c × KnowSum_c interaction indicates that toddlers’ similarity judgements converge more closely with the adult structure in colour regions for which the child has acquired the relevant lexical labels. Critically, this effect held after controlling for age, which argues against the interpretation that the relationship is driven solely by general cognitive maturation. This pattern is consistent with the weak Sapir–Whorf hypothesis as applied to colour ^27,28^: language does not construct the global colour-experience structure *de novo*, but rather refines local regions of the subjective colour space by sharpening the perceptual boundaries that lexical categories encode.

Our findings bear on the broader debate about the developmental origins of colour experience. The observation that the global relational structure of colour experience is largely shared between toddlers and adults aligns with the view that colour experience is primarily shaped by biologically constrained, prelinguistic mechanisms ^19,21,22^. At the same time, the misalignment associated with incomplete colour-word knowledge is consistent with evidence that linguistic categories can modulate colour processing at specific category boundaries ^27–29^. Importantly, the effects of language appear to operate primarily on a specific colour: light blue, rather than reorganising the global architecture. Recent evidence further supports this interpretation, demonstrating that basic colour-word comprehension produces a modest shift in categorical perception at category boundaries, but characterised this as a refinement of preexisting infant categories rather than category creation ^43^. Our data extend this emerging consensus from discrimination and memory paradigms to the domain of relational structures, suggesting that even at this higher-order level of experiential organisation, language serves a modulatory rather than constitutive role.

Several limitations should be noted. First, the toddler group provided binary (similar/dissimilar) rather than graded (4-point) similarity judgements, necessitated by the cognitive demands of the task. Although we confirmed the reliability of responses via double-pass correlations, the reduced resolution may have introduced noise or attenuated structural detail relative to the older groups. Second, the toddlers rated only a subset of colour pairs (30 out of 169), and group-level aggregation was required to construct the full dissimilarity matrix. This design precludes individual-level GWOT analysis for toddlers, and future work should seek methods that enable complete pairwise data collection within this age range. Third, our stimulus set comprised 13 colours chosen to correspond to the basic Japanese colour vocabulary; a larger, more nuanced stimulus set might reveal additional regions of alignment or misalignment that are not detectable at this resolution. Finally, the present design is cross-sectional, and longitudinal data would be needed to establish whether the observed relationship between colour-word acquisition and similarity-structure refinement reflects a causal influence or a shared developmental trajectory.

In conclusion, the application of unsupervised alignment via GWOT to developmental colour-experience data provides quantitative evidence that the global relational structure of colour experience is largely in place by toddlerhood, well before colour-term knowledge has fully stabilised. Language modulates local regions of this structure—particularly those where lexical boundaries are still being acquired—but does not appear to construct the overall geometry *de novo*. These findings bridge developmental psychology and consciousness science, offering a principled, label-independent framework for investigating how subjective experience emerges and transforms across development. The approach is inherently generalisable: by extending the similarity-structure paradigm to other modalities, cultures, and populations, future work can chart both the shared and idiosyncratic features of conscious experience across the human lifespan.

## Methods Ethics

This study was approved by the Ethics Committee of the Psychological Science Unit, Kyoto University (No. 3-P-22). Written informed consent was obtained from adult participants and from the parents of all child participants.

### Participants

We recruited 40 Japanese toddlers (mean age = 38.86 months, SD = 5.20; 26 girls), 32 preschool children (mean age = 66.13 months, SD = 13.71; 18 girls), and 32 adults (mean age = 31.56 years, SD = 3.66; 28 women) from a participant database. Sample sizes were determined by reference to previous developmental MDS work using comparable methods ^14^, which typically include approximately 30 participants per age group. Sex distributions differed across the three age groups, p = .006. Holm-corrected pairwise comparisons indicated a significant difference only between the preschool and adult groups (adjusted p = .004). However, because the colour structures of these two groups were perfectly aligned, this imbalance is unlikely to account for the present findings.

### Stimuli

The task was programmed in jsPsych. We used a set of 10 saturated colours with equi-hue spacing in HSV colour space: red, orange, yellow, light green, green, light blue, blue, purple, brown, and pink plus white, black, and grey. These colours correspond to the basic colour terms in Japanese ^39^. All stimuli were presented on a random noise background. The stimulus diameter was 140 px. In each trial, two stimuli were sequentially presented, positioned at two of 9 possible locations on an invisible circle (diameter 500 px, centred on the screen, with locations spaced at 40° increments from 0° to 320°) to ensure equal eccentricity and prevent overlap with screen edges and response buttons.

## Procedure

### Colour similarity task

After consent, participants rated the similarity of two sequentially presented colour patches. The toddler group used a 2-point scale (similar/dissimilar); the preschool children and adult groups used a 4-point scale (very dissimilar, slightly dissimilar, slightly similar, very similar). The main experiment comprised 30 trials (a subset of the 169 pairs) for toddlers and 169 trials (all 13 × 13 ordered colour-pair trials, including same-colour pairs) for preschool children and adults. To assess response reliability, the first 10 trials (toddlers) or 33 trials (older groups) were repeated, yielding 40 and 202 trials in total (double-pass design). Trial order and stimulus locations were fixed across participants in the preschool and adult groups, whereas toddlers received different subsets of colour pairs. Each trial began with a central fixation cross (500 ms), followed by one colour patch (500 ms) and then by another (500 ms), and then the response screen.

### Colour-word tasks

For the toddler group, two additional tasks assessed colour-term knowledge about 13 colours in the colour similarity task. In the comprehension task, children pointed to a named colour from among an array. In the labelling task, children were shown a colour patch and asked to name it. Performance was scored as the proportion of correct responses (0–1 per colour per task).

## Data analysis

### Exclusion criteria

Participants were excluded if they did not complete the main experiment or if their double-pass Pearson correlation fell below 0.30 ^35^. After exclusions, 31 toddlers, 28 children, and 32 adults were retained.

### Dissimilarity matrices

For preschool children and adults, individual dissimilarity matrices were averaged to produce group-level matrices. For toddlers, individual partial matrices were integrated into a single group-level matrix by averaging available values for each colour pair.

*Representational similarity analysis.* Upper-triangular values of each group-level dissimilarity matrix were extracted, and pairwise Spearman rank correlations were computed. Statistical significance was assessed via permutation-based Mantel tests (5,000 permutations, two-sided).

### Gromov–Wasserstein optimal transport (GWOT)

Colour similarity structures were estimated by applying MDS to the group-level dissimilarity matrices^7,16^. Next, we performed GWOT to identify the optimal transportation plan Γ between each pair of structures ^7,16^. The alignment was evaluated using the top-*k* matching rate, defined as the proportion of colours for which the correct physical counterpart fell within the top-*k* mappings under Γ. To search for the global optimum, we performed 200 optimisation iterations with different ε values (controlling entropy regularisation) and random initialisations. The solution with the lowest GW distance was selected.

### Leave-one-colour-out analysis

To identify localised sources of misalignment, we iteratively removed each of the 13 colours and recomputed the GWOT alignment on the remaining 12 colours. *GLMM.* We fit a GLMM with a logit link (lme4::glmer) predicting children’s binary similarity response from: (a) the adult similarity anchor (Adult_c; group-mean adult dissimilarity, reverse-scored and mean-centred), (b) a trial-level colour-word knowledge score (KnowSum_c; sum of comprehension and labelling scores for the two colours, 0–4, mean-centred), (c) their interaction, and (d) mean-centred age (age_c). Random intercepts were included for participant and colour pair. All analyses were performed in R (version 4.2.3).

## Data availability

All experimental data and task materials are available at https://github.com/RyoichiWata/Color_and_Label_2026/tree/paper_minimal_data.

## Acknowledgements

Y.M., N.S., and M.O. were supported by a Grant-in-Aid for Transformative Research Areas (23H04832, 23H04835, 23H04834) from the Japan Society for the Promotion of Science. The funders had no role in study design, data collection and analysis, decision to publish, or manuscript preparation.

## Author contributions

R.W., N.S., and Y.M. developed the study concept and collected the data. R.W., M.O., and Y.M. performed the analyses, and J.W. and M.O. created the visualisations. R.W. and Y.M. drafted the manuscript, and J.W., M.O., and N.S. revised the manuscript. All authors approved the final manuscript for submission.

## References

1. Locke, J. An Essay Concerning Human Understanding. (Clarendon Press, Oxford, 1689).

2. Palmer, S. E. On qualia, relations, and structure in color experience. Behav Brain Sci 22, 976–985 (1999).

3. Shoemaker, S. The inverted spectrum. J. Philos. 79, 357 (1982).

4. Dennett, D. C. Quining qualia. In Consciousness in contemporary science (eds. A. J. Marcel & E. Bisiach) 42–77 (Clarendon Press/Oxford University Press., 1988).

5. Kanai, R. & Tsuchiya, N. Qualia. Curr. Biol. 22, R392–6 (2012).

6. Nagel, T. What Is It Like to Be a Bat. The Philosophical Review 83, 435–450 (1974).

7. Kawakita, G., Zeleznikow-Johnston, A., Takeda, K., Tsuchiya, N. & Oizumi, M. Is my “red” your “red”?: Evaluating structural correspondences between color similarity judgments using unsupervised alignment. iScience 28, 112029 (2025).

8. Moriguchi, Y., Tsuchiya, N. & Saigo, H. A structural approach in cognitive developmental research with category theory. Human Development (2026).

9. Tsuchiya, N. & Saigo, H. A relational approach to consciousness: Categories of level and contents of consciousness. Neurosci Conscious 2021, niab034 (2021).

10. Shepard, R. N. & Cooper, L. A. Representation of colors in the blind, color-blind, and normally sighted. Psychol. Sci. 3, 97–104 (1992).

11. Kleiner, J. Towards a structural turn in consciousness science. Conscious. Cogn. 119, 103653 (2024).

12. Helm, C. E. Multidimensional ratio scaling analysis of perceived color relations. J. Opt. Soc. Am. 54, 256–262 (1964).

13. Zeleznikow-Johnston, A., Aizawa, Y., Yamada, M. & Tsuchiya, N. Are color experiences the same across the visual field? J. Cogn. Neurosci. 35, 509–542 (2023).

14. Moriguchi, Y. et al. Comparing color qualia structures through a novel similarity task in young children versus adults. Proc. Natl. Acad. Sci. U. S. A. 122, e2415346122 (2025).

15. Kriegeskorte, N., Mur, M. & Bandettini, P. Representational similarity analysis - connecting the branches of systems neuroscience. Front. Syst. Neurosci. 2, 4 (2008).

16. Togashi, Y., Yotsumoto, Y., Hiramatsu, C., Tsuchiya, N. & Oizumi, M. Robust individual alignment of color qualia structures: Toward a structure-based taxonomy of divergent color experiences. bioRxiv (2026) doi:10.64898/2026.02.13.705699.

17. Mémoli, F. Gromov–Wasserstein distances and the metric approach to object matching. Found. Comput. Math. 11, 417–487 (2011).

18. Yang, J., Kanazawa, S., Yamaguchi, M. K. & Kuriki, I. Cortical response to categorical color perception in infants investigated by near-infrared spectroscopy. Proceedings of the National Academy of Sciences 113, 2370–2375 (2016).

19. Bornstein, M. H., Kessen, W. & Weiskopf, S. Color vision and hue categorization in young human infants. J. Exp. Psychol. Hum. Percept. Perform. 2, 115–129 (1976).

20. Franklin, A. et al. Salience of primary and secondary colours in infancy. British Journal of Developmental Psychology 26, 471–483 (2008).

21. Maule, J., Skelton, A. E. & Franklin, A. The development of color perception and cognition. Annu. Rev. Psychol. 74, 87–111 (2023).

22. Skelton, A. E., Catchpole, G., Abbott, J. T., Bosten, J. M. & Franklin, A. Biological origins of color categorization. Proc. Natl. Acad. Sci. U. S. A. 114, 5545–5550 (2017).

23. Özgen, E. Language, learning, and color perception. Curr. Dir. Psychol. Sci. 13, 95–98 (2004).

24. Witzel, C. & Gegenfurtner, K. R. Is there a lateralized category effect for color? J. Vis. 11, 16 (2011).

25. Forder, L. & Lupyan, G. Hearing words changes color perception: Facilitation of color discrimination by verbal and visual cues. J. Exp. Psychol. Gen. 148, 1105–1123 (2019).

26. Goldstein, J., Davidoff, J. & Roberson, D. Knowing color terms enhances recognition: Further evidence from English and Himba. Journal of experimental child psychology 102, 219–238 (2009).

27. Gilbert, A. L., Regier, T., Kay, P. & Ivry, R. B. Whorf hypothesis is supported in the right visual field but not the left. Proc. Natl. Acad. Sci. U. S. A. 103, 489–494 (2006).

28. Winawer, J. et al. Russian blues reveal effects of language on color discrimination. Proc. Natl. Acad. Sci. U. S. A. 104, 7780–7785 (2007).

29. Franklin, A. et al. Lateralization of categorical perception of color changes with color term acquisition. Proc. Natl. Acad. Sci. U. S. A. 105, 18221–18225 (2008).

30. Pitchford, N. & Mullen, K. The development of conceptual colour categories in pre-school children: Influence of perceptual categorization. Vis. cogn. 10, 51–77 (2003).

31. Wagner, K., Dobkins, K. & Barner, D. Slow mapping: Color word learning as a gradual inductive process. Cognition 127, 307–317 (2013).

32. Saji, N., Imai, M. & Asano, M. Acquisition of the meaning of the word orange requires understanding of the meanings of red, pink, and purple: Constructing a lexicon as a connected system. Cogn. Sci. 44, e12813 (2020).

33. Pitchford, N. J. & Mullen, K. T. Is the acquisition of basic-colour terms in young children constrained? Perception 31, 1349–1370 (2002).

34. Forbes, S. H. & Plunkett, K. Linguistic and cultural variation in early color word learning. Child Dev. 91, 28–42 (2020).

35. Watanabe, R. & Moriguchi, Y. The double-pass correlation: A reliability metric for subjective judgments in developmental research. MethodsX 17, 104039 (2026).

36. Chambers, C. T. & Johnston, C. Developmental differences in children’s use of rating scales. J. Pediatr. Psychol. 27, 27–36 (2002).

37. Mellor, D. & Moore, K. The use of Likert scales with children. J. Pediatr. Psychol. 39, 369–379 (2014).

38. Kong, L., Li, J., Tang, J. & So, A. M.-C. Outlier-Robust Gromov-Wasserstein for Graph Data. In Advances in Neural Information Processing Systems 36 (eds. Oh, A. et al.) vol. 36 24781–24803 (Neural Information Processing Systems Foundation, Inc. (NeurIPS), San Diego, California, USA, 2023).

39. Kuriki, I. et al. The modern Japanese color lexicon. J. Vis. 17, 1 (2017).

40. Uchikawa, K. & Boynton, R. M. Categorical color perception of Japanese observers: Comparison with that of Americans. Vision Res. 27, 1825–1833 (1987).

41. Beekhuizen, B. & Stevenson, S. More than the eye can see: A computational model of color term acquisition and color discrimination. Cogn. Sci. 42, 2699–2734 (2018).

42. Yang, J. et al. Slow mapping in color word acquisition across languages: Evidence from Japanese children. *Front*. Dev. Psychol. 3, (2025).

43. Forbes, S. H. & Plunkett, K. Colour perception changes with basic colour word comprehension. Dev. Sci. 26, e13406 (2023).

