## Supplemental Figures for "Unsupervised alignment reveals shared but locally refined colour qualia structure from toddlerhood to adulthood"

Supplementary Information

Supplementary Figure S1-S4

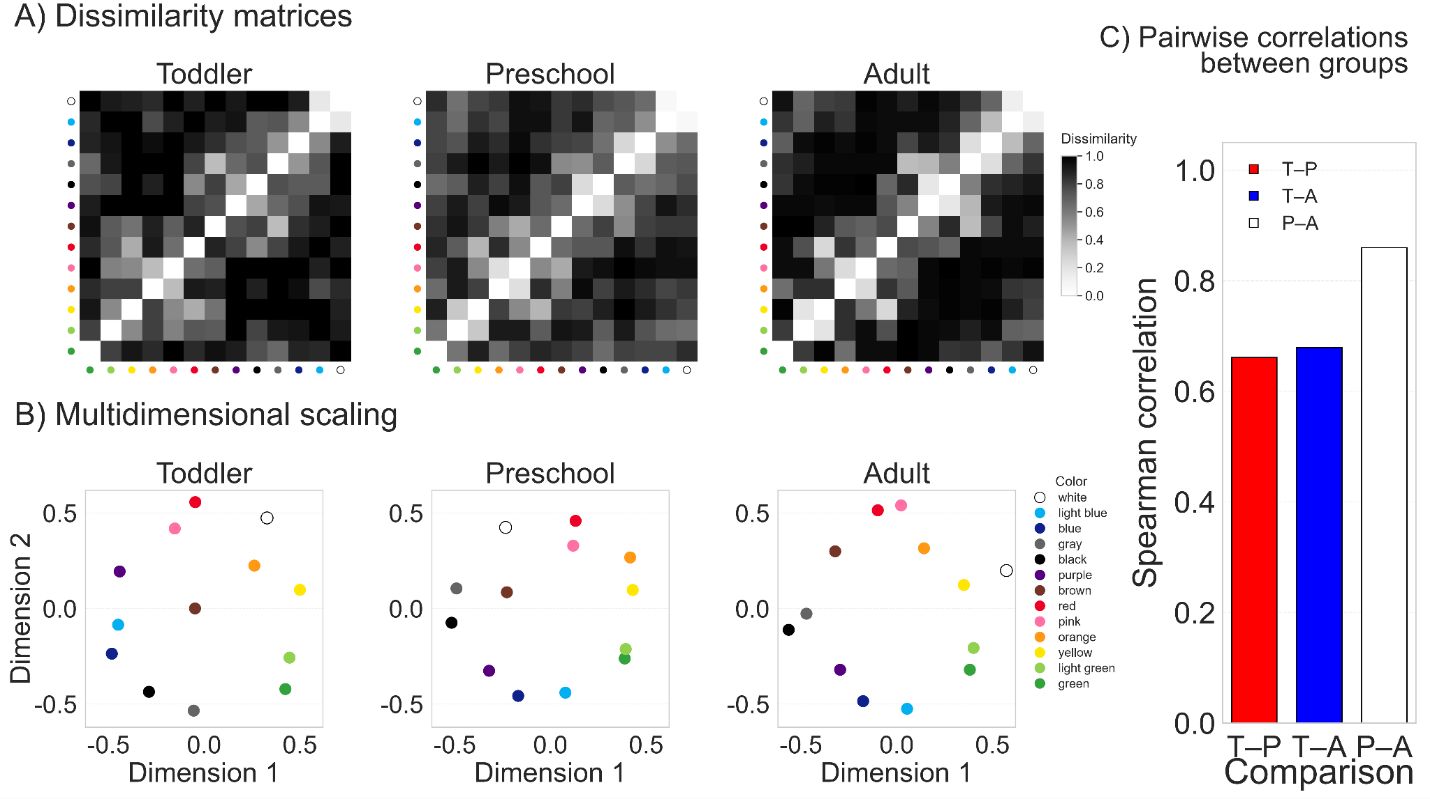

Figure S1. Conventional RSA based on binary-recoded similarity ratings. A) Group-mean dissimilarity matrices for each age group. The 4-point ratings obtained from preschoolers and adults were recoded into a binary scale corresponding to the toddler response format. White and dark cells indicate greater similarity and greater dissimilarity, respectively. B) Two-dimensional MDS representations for each age group. Symbol colors correspond to the stimulus colors used in the experiment. C) Pairwise correlations between the group-mean dissimilarity matrices. T–P, T–A, and P–A indicate toddler–preschooler, toddler–adult, and preschooler–adult comparisons, respectively.

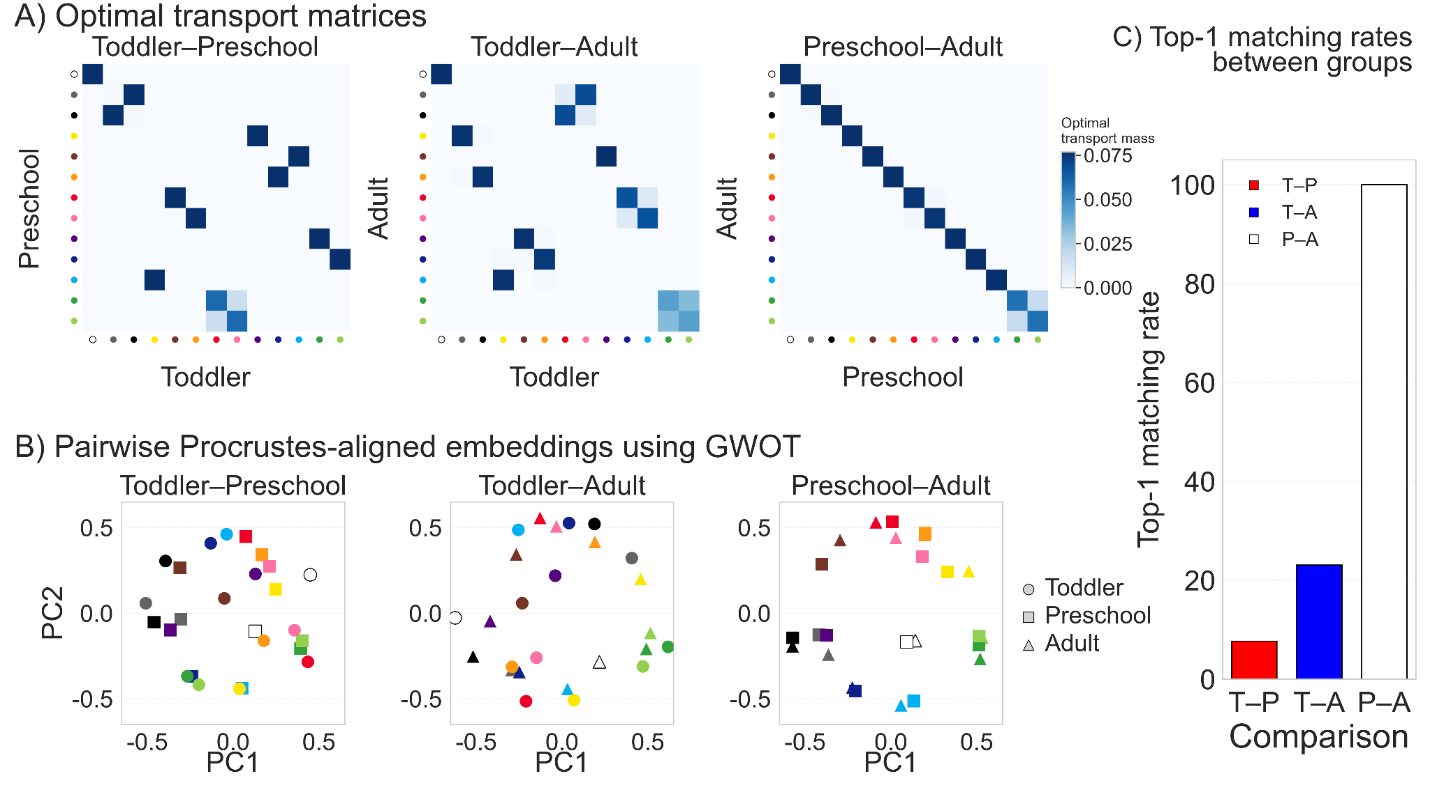

Figure S2. Analyses using Gromov–Wasserstein optimal transport (GWOT) with binary-recoded similarity ratings. A) Optimal transport matrices between age groups. For preschoolers and adults, the original 4-point similarity ratings were recoded into a binary scale corresponding to the toddler response format. B) Pairwise Procrustes-aligned embeddings based on the GWOT correspondence between groups. C) Top-1 matching rates between groups. T–P indicates the toddler–preschooler comparison, T–A indicates the toddler–adult comparison, and P–A indicates the preschooler–adult comparison.

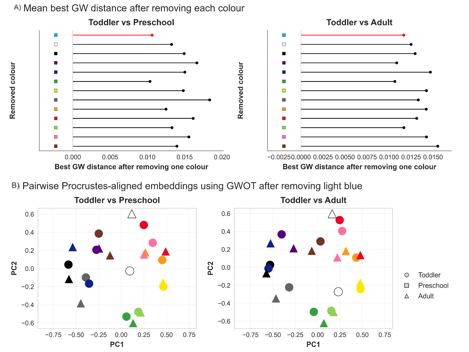

Figure S3. A) Mean best GW distance after removing each colour. B) Pairwise Procrustes-aligned embeddings using GWOT after removing light blue. Colour-specific relational-profile similarity between toddlers and older groups.

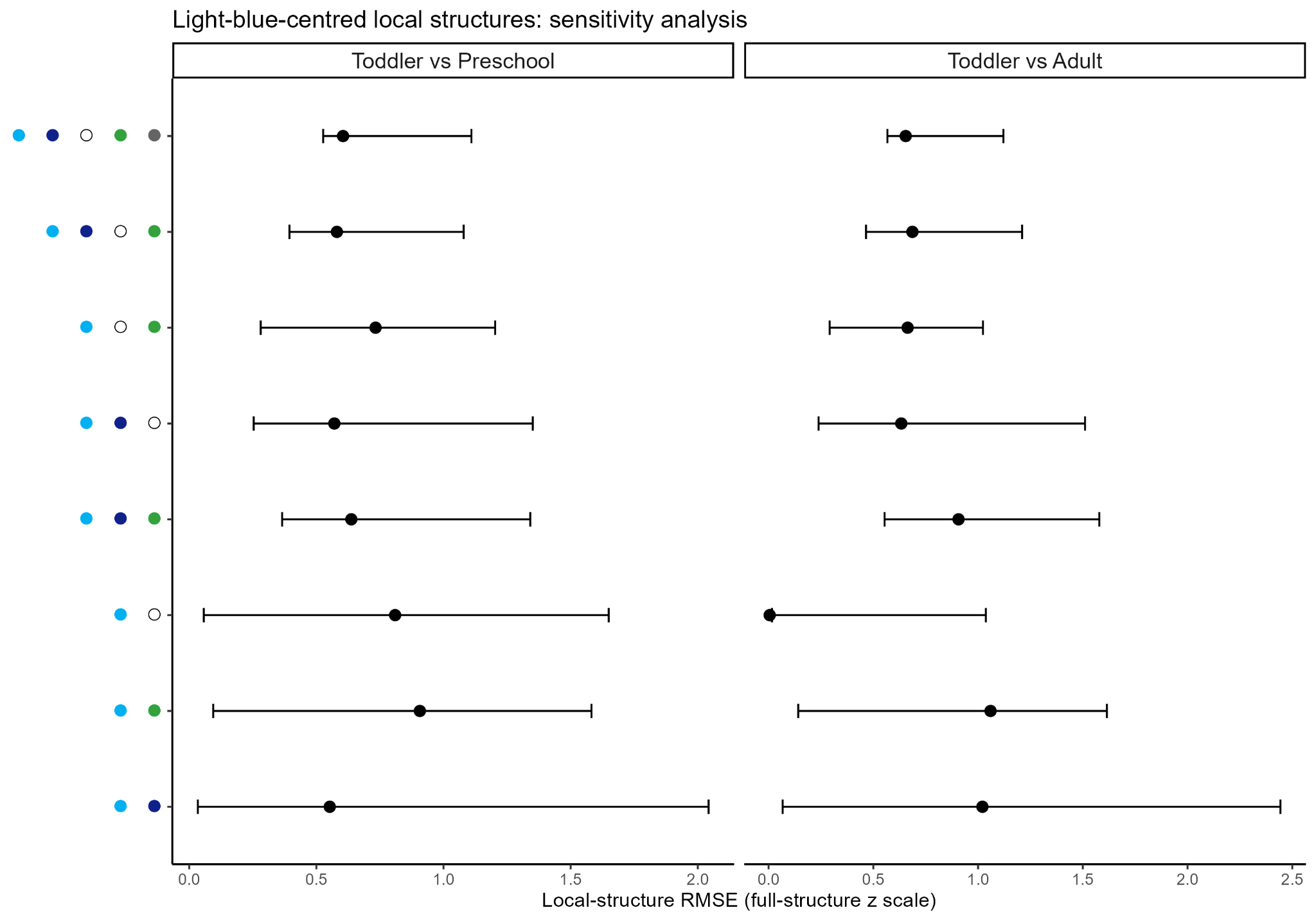

Figure S4. Local-structure discrepancy was quantified from the six within-subset pairwise dissimilarities among light blue, blue, white, and green. Although this light-blue-centred subset showed some developmental discrepancy, it was not reliably more discrepant than equivalently sized subsets that did not contain light blue. Sensitivity analyses using light-blue-centred subsets containing two to five colours showed the same general pattern, with no contrast intervals excluding zero.
